# *Lactobacillus* maintains IFNγ homeostasis to promote behavioral stress resilience

**DOI:** 10.1101/2023.05.10.540223

**Authors:** Andrea R. Merchak, Samuel Wachamo, Lucille C. Brown, Alisha Thakur, Brett Moreau, Ryan M. Brown, Courtney Rivet-Noor, Tula Raghavan, Alban Gaultier

## Abstract

The gut microbiome consists of the trillions of bacteria, fungi, and viruses that inhabit the digestive tract. These communities are sensitive to disruption from environmental exposures ranging from diet changes to illness. Disruption of the community of lactic acid producing bacteria, *Lactobaccillacea*, has been well documented in mood disorders and stress exposure. In fact, oral supplement with many *Lactobacillus* species can ameliorate these effects, preventing depression- and anxiety-like behavior. Here, for the first time, we utilize a gnotobiotic mouse colonized with the Altered Schaedler Flora to remove the two native species of *Lactobaccillacea*. Using this novel microbial community, we found that the *Lactobacillus* species themselves, and not the disrupted microbial communities are protective from environmental stressors. Further, we determine that *Lactobaccillacea* are maintaining homeostatic IFNγ levels which are mediating these behavioral and circuit level responses. By utilizing the Altered Schaedler Flora, we have gained new insight into how probiotics influence behavior and give novel methods to study potential therapies developed to treat mood disorders.

## Introduction

The gut microbiota consists of a collection of bacteria, viruses, and fungi that maintains symbiotic homeostasis within the digestive tract. This community is important for proper immune development, digestion, as well as many other functions[1,2]. The connection between the gut microbiota and the brain has re-emerged as an area of focus for biomedical research related to psychological disorders[3–6].

Dysbiosis, or a disruption of the microbial community, is commonly reported in patients with psychological stress or mood disorders[7,8]. Numerous studies have found *Lactobacillus* to be one of the primary bacterial families diminished [5,9–11] in such disorders. In fact, in clinical trials and animal preclinical research, *Lactobacillus* was found to be a psychobiotic which aids in stress resistance, reduces disordered behavior in mice, and reduces both self-reported depression and anxiety in patients [5,12–16]. Interestingly, the beneficial effects of *Lactobacillus* in our gut has been observed using different species and strains of *Lactobacillus*, suggesting that this is a generalized phenomenon to the bacterial family *Lactobacillacea*[17]. Due to the difficulty of executing gnotobiotic studies in specific pathogen free (SPF) mice and the fact that *Lactobacilli* are commensal flora, almost all work thus far has been done in an additive manner via oral supplement. There is a fundamental lack of understanding of how a complete loss of *Lactobacilli* could affect host physiology.

To circumvent these limitations, this study capitalizes on the underutilized gnotobiotic consortia known as the Altered Schaedler Flora (ASF). This bacterial consortium was standardized in 1978 and consists of eight well-characterized bacterial strains passed from mother to pup during birth and development [18]. These bacteria include two strains of *Lactobacillus*, two strains of *Clostridium*, and one strain of each of *Bacteroides, Mucispirillum, Eubacterium and Pseudoflavonifactor*. We harnessed this powerful tool, and the two species of *Lactobacillus* were removed to better understand their role in behavior, immune development, and mood.

By utilizing mice lacking *Lactobacilli* from birth, we found that Type 1 adaptive immunity is important as a primary mediating factor in stress resistance. Building off a body of work showing that *Lactobacillus* is necessary for the maintenance of systemic IFNγ, we further demonstrate that both *Lactobacilli* and IFNγ are necessary for resilience to environmental stressors.

## 1. Materials and Methods

### Mice

C57BL/6J (#000664) were purchased from Jackson Laboratory. Germ free mice (B6NTac), ASF(+L), and ASF(-L) mice were purchased from Taconic and maintained in Flexible Film Isolators (Class Biologically Clean) with a monthly QC to confirm germ status. Microbiomes were validated by Taconic, upon arrival, and periodically in-house. Mice were sterile-transferred to Semi-Rigid Isolators (PARKBIO Inc.) for all experimental manipulations. All mice had a 12-hour light/dark schedule. All procedures were approved by the University of Virginia ACUC (protocol #1918).

### Chronic Restraint Stress

For the duration of the Chronic Restraint stress (CRS), mice were housed individually. Mice were exposed to two hours of daily restraint at a different time of day. Additionally, they were exposed to one of three random overnight stressors that included moist bedding, tilted cage, or two cage changes in a 24 hour period [5,19]. After three weeks of stress, behavioral assays were performed and fecal collection began for microbiome transfer. The CRS and fecal collection were continued for another two weeks.

### Microbiome Transfer

For two weeks, dirty bedding from each group was transferred daily to the cages of female germ free C57BL/6J mice. Two weeks passed before the ex-germ-free animals were used for experiments.

### Acute and Subclinical Stress

Mice were housed in sterile semi-rigid isolators. For the mild acute stressor, mice were removed from the isolator, placed in a biological safety cabinet, and restrained in 50mL conical vials with ventilation holes for three hours. During this time, naïve controls remained in their home cages in the semi-rigid isolator. Approximately 30 min-1 hour after stress exposure, the mice were euthanized and brains were collected for c-Fos staining. For the subclinical chronic stress, autoclaved 50mL conical vials with ventilation holes were passed into the semi-rigid isolators. Mice were restrained in the conical vials for two hours daily, at a different time of day, for seven days. On the eighth day, behavioral testing began.

### c-Fos Staining

Mice were perfused with saline containing heparin (10,000U/L; Sigma; H-3125) followed by PBS containing 4% paraformaldehyde. Brains were dissected and drop fixed in PBS paraformaldehyde for 24 hours, at which point they were transferred to 30% sucrose solution and mounted in cryomolds with OCT (Tissue-Tek; 4583). Cryosections (30μm thickness) were stained with rabbit anti-c-Fos (abcam; ab190289) and secondary AlexaFluor 488 tagged anti-Rabbit (Life Technologies; A21206). Sections were mounted using Prolong Gold Anti-Fade Reagent (Life Technologies, P36980) and imaged with a Leica TCS SP8 confocal microscope. Images were analyzed and c-Fos positive cells were counted by a blinded researcher using unbiased thresholding using Imaris version 9.9.1.

### Behavioral Tests

Prior to all behavioral tests, animals were acclimated to the room for 1 hour in a biological safety cabinet. Animals were kept in the biological safety cabinet until testing, after which they were returned to the biological safety cabinet. All tools were cleaned before testing with MinnCare Disinfectant. Between mice, tools were cleaned with 70% ethanol and between groups, tools were cleaned again with MinnCare Disinfectant.

#### Tail Suspension

Mice were taped by the tip of a tail to a surface two feet high for six minutes. Two minutes were used for acclimation and only the last four minutes were used for analysis. Video was recorded on a Hero Session 5 GoPro and analyzed with Noldus behavioral analysis software.

#### Nestlet Shred

Animals were placed into newly autoclaved cages with corncob bedding and allowed to acclimate for 30 minutes. A pre-weighed nestlet was placed in the center of the cage and left for 30 minutes. The nestlet was removed from the cage and weighed again.

#### Elevated Plus Maze

Mice were placed in the center of an elevated plus maze and tracked using video recorded from a Hero Session 5 GoPro for ten minutes. The amount of time spent in the open arms was calculated using Noldus behavioral analysis software.

### 16S sequencing

16S Sequencing was conducted by Microbiome Insights. Initial sequence data was analyzed using the latest version of Quantitative Insights Into Microbial Ecology 2 (Qiime2 v2021.11)[20]. Demultiplexed paired-end sequence reads were preprocessed using DADA2[21]. The first 20 base pairs were trimmed from forward and reverse reads before they were merged to remove adaptors. Taxonomy was assigned to Amplicon Sequence Variants (ASVs) using a Naïve Bayes classifier trained on full length 16S sequences from the latest Greengenes16S database (13_8) clustered at 99% sequence similarity. Samples were rarified before core diversity analysis. Core diversity metrics were analyzed, including number of ASVs. Nonmetric multidimensional scaling was performed in RStudio using the phyloseq package[22].

### Serum Collection and Luminex

Blood was collected by cardiac puncture and transferred to BD Microtainer SST(BD; 365967) to prepare serum per manufacturer’s instructions. Samples were frozen at -80°C until downstream analysis. Mouse Luminex panel was run per manufacturer’s instructions on a MagPix by the University of Virginia Flow Cytometry Core, RRID: DCR_0177829.

### Untargeted Metabolomics

Untargeted Metabolomics was performed by the University of Michigan Biomedical Research Core Facilities. Serum was collected by cardiac puncture as described before and frozen at -80 °C. Data was annotated using *Binner*[23]. Data was pre-processed for analysis and features that discriminate between groups were identified using Sparse Partial Least Squares Discriminant Analysis (sPLS-DA) with component set at one and selecting 50 variables.

### RNA Extraction and Quantitative PCR

After Euthasol injection, mice were perfused with saline with heparin (10,000U/L; Sigma; H-3125). The duodenum was dissected and 0.5 cm sections were used for RNA extraction. Extraction was conducted using the Bioline Isolate II RNA mini kit per manufacture’s protocol (BIO-52073). RNA was normalized after quantification using the Biotek Epoch Microplate Spectrophotometer and reverse transcribed using the Bioline SensiFast cDNA Synthesis Kit (BIO-65054). Primers for qPCR are listed in the supplemental materials. Results were analyzed using the relative quantity (ΔΔcq) method.

### Flow Cytometry

Animals were perfused with saline with heparin (10,000U/L; Sigma; H-3125). Peyer’s patches, lymph nodes, and spleens were mashed through 70 μm filters to achieve single-cell suspensions [24].

Dural meninges were dissected from skull caps into RPMI. They were then added to 2 mL of digestion buffer (RPMI containing Collagenase D (2mg/mL; Sigma-Aldrich; 11088882001), Collagenase 11 (2mg/mL; Gibco; 17101-015), DNAse (20U/mL; Worthington; LS002139)) in a 15mL conical vial. They were agitated at 37°C for 30 minutes. 20μL of 0.5M EDTA was then added and filtered through a 70μm filter. 2mL RPMI with 5% FBS was added followed by 5 minutes of centrifugation at 400g. The cells were subsequently resuspended in FACS buffer. Small intestines (duodenum, jejunum, and ilium) were dissected, flayed open, and cut into 1cm long pieces. These pieces were washed two times in 30mL of HBSS-/-with 5% FBS via agitation at 37°C for 20 minutes. After each wash, excess liquid was removed by pouring the tissue over a mosquito net. After washing, the tissue was minced and added to a new 50mL conical vial with 20mL of digestion buffer (HBSS-/-, 5% FBS), Collagenase 8 (900U/mL; Milipore; C2139), and DNAse (20U/mL; Worthington; LS002139). These were agitated at 37°C for 30 minutes then vigorously vortexed and filtered through 70μm filter. 30mL of HBSS-/-with 5% FBS was added to halt digestion and centrifuged at 400g for 5 minutes. The pellet was resuspended in 50mL of HBSS-/-with 5% FBS and centrifuged again at 400g for 5 minutes. Finally, the pellet was resuspended in 1mL of FACS and 200μL was used for flow staining.

For staining, single cell suspensions were incubated with Fc Block CD16/32 (Invitrogen; 14-0161-85). The cells were then incubated for 30 minutes with the surface antibodies and Live/Dead Ghost Dye Violet 510 (Tonbow Biosciences; 13-0870). Cells were washed and the FoxP3/Transcription Factor Staining Kit (eBioscience; 00-5523-00) was used per manufacturer’s instructions in order to do intracellular staining. OneComp eBeads (Thermo Fisher Scientific; 01-111-42) were used for single color controls. Flow cytometry was performed on a Beckman Coulter Gallios flow cytometer and the data were analyzed with FlowJo software v10.7.1.

### Modified LB media

Modified LB media consisted of LB Base (Sigma, L3022-250G) supplemented with 0.25g/L L-cysteine (filter sterilized; Sigma, 30120-10G). Prior to culture, the media was sterilized and then supplemented with 10 mL/L hemin solution (filter sterilized 0.5 g/L dissolved in 1% NaOH, 99% deionized water; Sigma H9030-1G), 100μL/L vitamin K1 solution (0.5% vitamin K1 dissolved in 95% ethanol; Sigma, V3501-1G), 10mL/L lactose solution (filter sterilized5 mg/mL dissolved in deionized water; cat), 10mL/L Tween 20 (filter sterilized; 1mg/mL dissolved in deionized water), 39mL/L of a mineral salts solution (containing 6g/L KH2PO4, 6g/L (NH4)2SO4, 12g/L NaCl, 2.5g/L MgSO4,7H2O, 1.6 g/L CaCl2,2H2O), all dissolved in deionized water and filter sterilized).

### Lactobacillus EV isolation

Whole intestinal contents (small intestine, cecum, and colon) from two pooled ASF or ASF-L mice were sterile harvested and passed into an anaerobic chamber. Under anaerobic conditions, the intestines were flayed open and shaken in modified LB media. Solids were removed using a sterile 70 μm filter and cultured, shaking for 36 hours at 37°C. Bacterial cultures were removed from the anaerobic chamber and pelleted by centrifugation at 10,000g for 20 min. Supernatant was passed through a 0.22μm filter. Filtrate was concentrated using 10KDa concentrators (Amicon; UFC901024) by spinning at 3000g at 4°C until the original 250mL was reduced to around 15mL. Extracellular vesicles were pelleted by ultracentrifugation at 150,000g for 3 hours at 4°C. Pellets were resuspended in filtered PBS and centrifuged again at 150,000g for 3 hours at 4°C. Pellet was resuspended in 550μL filtered PBS, protein was quantified using a BCA assay and samples were stored at -80°C until use.

### Isolation and Preparation of T cells

Mesenteric, inguinal, axial, and brachial lymph nodes plus the spleen were dissected from the C57/BL6 mice after CO_2_ euthanasia. They were then dissociated through sterile 70 μm filters and lysed with ACK lysis buffer (Quality Biological, 118-156-101). An EasySep Mouse CD4+ T cell Isolation Kit (Stem Cell Technologies, #19852) was used per manufacturer’s protocol to sort naïve CD4+ T cells. The cells were incubated at 1×10 cells/mL in skew media in anti-CD28 (InVivoMab, BE0015-5) and anti-CD3 (InVivoMab, BE0001-1) coated plates as previously described[25]. T_H_1 and T_H_2 media contained RPMI (Gibco, 11875-093) supplemented with 10% FBS (Optima, S12450H), 0.5% Pen/Strep (Gibco, 15240-062), 1mM sodium pyruvate (Gibco, 11360-070), 2mM L-glutamine (Gibco, 25030081), 10mM HEPES (Gibco, 15603-080), 1:100 NEAA (Gibco, 11140-050), and 50uM Β-mercaptoethanol (Fisher Scientific, O3446I-100). Treg and T_H_17 media contained IMDM (Gibco, 12440-053) using the same supplements. T_H_1 skew media contained IL-2 (100U/mL; Tecin, 23-6019), IL-12 (10ng/mL; PeproTech, 210-12), and anti-IL-4 (10ug/mL; InVivoMab, BE0045). T_H_2 skew media contained IL-2 (100U/mL; Tecin, 23-6019), IL-4 (10ng/mL; InVivoMab, BE0045), and anti-IFNγ (10ug/mL; InVivoMab BE0054). Treg skew media contained TGF-Β (5ng/mL; BioLegend, 763102) and anti-CD28 (2ug/mL; InVivoMab, BE0015-5). T_H_17 skew media contained IL-6 (20ng/mL; BioLegend, 575704), IL-23 (10ng/mL; Invitrogen, 14-8231), TGF-Β (0.3ng/mL; BioLegend, 763102), anti-IL-4 (10ug/mL; InVivoMab, BE0045), and anti-IFNγ (10ug/mL; InVivoMab BE0054). T_H_1 and T_H_2 cell cultures were expanded on day 3 after plating by transferring cells to larger plates and adding 3x volume of RPMI supplemented media containing IL-2 (100U/mL). On day 5, skewing was complete and cells were washed and plated for downstream assays. Treg and T_H_17 cell cultures were not expanded but simply washed and plated for downstream assays on day 4 after plating.

### ELISA

ELISA for IFNγ was performed as previously described [26]. Antibodies used were as follows: anti-IFNγ (BioLegend; 517902) and biotin labeled anti-IFNγ (BioLegend; 505704).

## 3.1 Results

### The transfer of microbiota from stressed mice to germ-free (GF) mice is sufficient to induce depressive- and anxiety-like behaviors

Microbiota changes are well documented in both mice and humans exposed to environmental stress [9,27–29]. Here, we used a previously characterized unpredictable chronic mild stress (UCRS) paradigm to induce depressive- and anxiety-like behaviors[5,19]. Mice were exposed to two randomized mild stressors each day for a period of 3 weeks. As previously reported, stressed mice spend more time inactive than naïve mice in the tail suspension test (**Figure 1A**) and had more repetitive behavior in the nestlet shred test (**Figure 1B**)[5,19]. Furthermore, these mice presented with microbiome dysbiosis (**Figure 1C-E**) and lower levels of *Lactobacillus* as measured by 16S sequencing when compared to naïve mice, replicating our previously published results [5]. There was no distinct group of bacteria that replaced the ecological niche created by the decrease of *Lactobacillus* (**Figure 1D**). To explore if microbiome dysbiosis is a marker of or an active contributor to anxiety- and depression-like behaviors, the microbiota from naïve and stressed animals was transferred to germ-free mice. To this end, we transferred bedding from the donor cages to the germ-free cages daily for two weeks. The mice were then left for two weeks to allow the microbiota to fully engraft before behavioral testing (**Figure 1F**). Strikingly, the depressive- and anxiety-like behaviors transferred to the ex-germ-free mice. The mice that received bedding from stressed mice spent more time inactive during tail suspension than the ex-germ-free mice housed with bedding obtained from naïve animals (**Figure 1G**). Mice colonized with the microbiome from stressed mice also presented with more anxiety-like behaviors than the naive recipients as measured with the elevated plus maze (**Figure 1H, I**). These data indicate that bacterial transfer is sufficient to drive behaviors associated with mood disorders and environmental stress exposure and these data support a growing body of evidence showing that the microbiota can directly modulate behavior [5,28,30–34].

**Figure 1:**
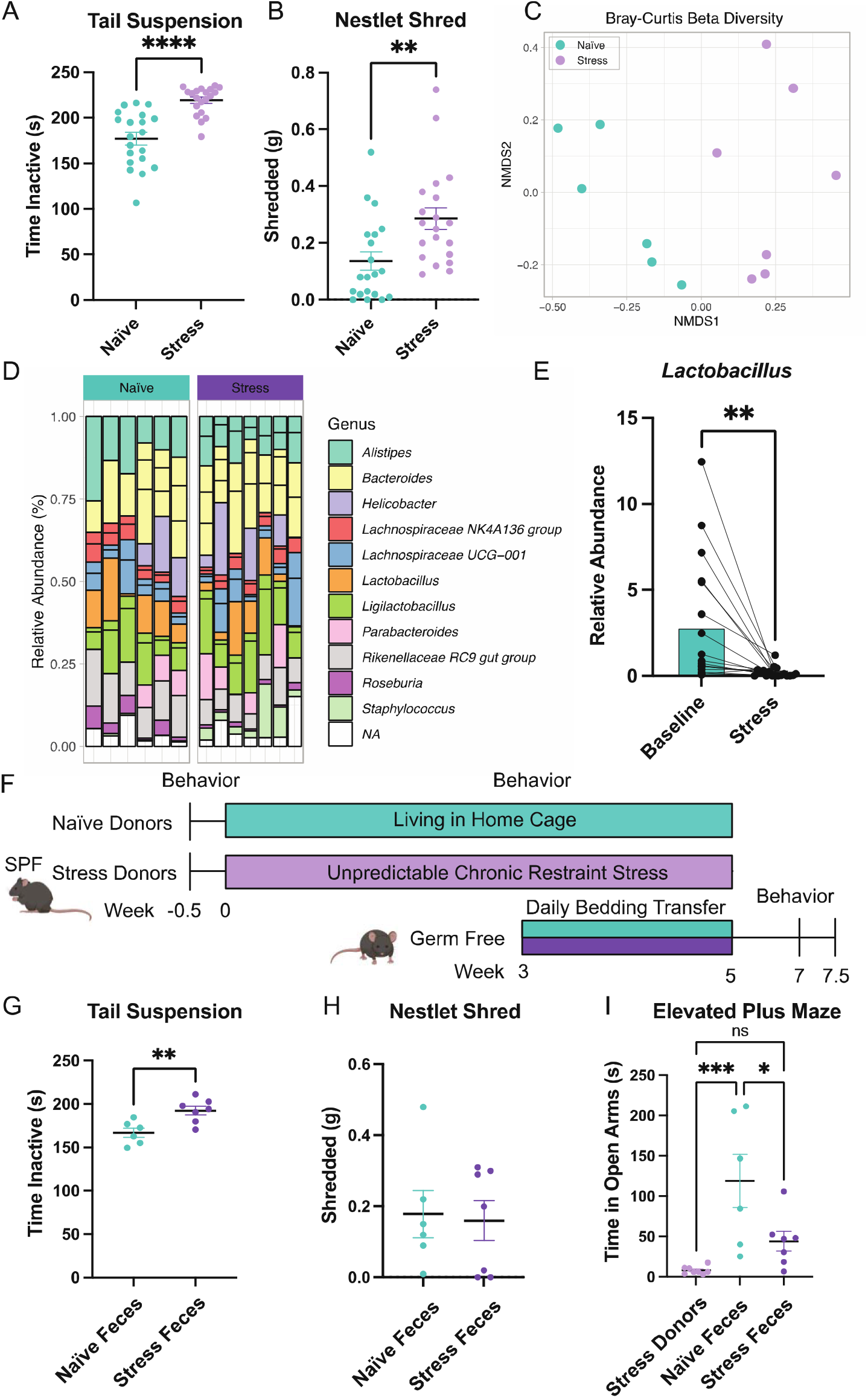
Gut microbiota drives anxiety- and depressive-like behavior. SPF C57BL/6J mice underwent the Unpredictable chronic mild stress (UCMS) paradigm. After three weeks of stress, behavior was performed and mice had increased (**A**) time inactive in the tail suspension test and (**B**) increased nestlet shred (Male mice; n=20 mice/group; N= 1 experimental replicate; Student’s T tests). These mice also had dysbiosis as measured by 16S sequencing. (**C**) The microbiome of stressed and naïve mice clustered separately by Bray-Curtis analysis. (**D**) There were no significantly changed genera of bacteria except for (E) *Lactobacillus* which was confirmed with qPCR (Male mice; n=6-7 mice/group; N= 1 experimental replicate; Student’s T tests). The stressed SPF mice and naïve mice served as fecal donors for germ-free mice. (**F**) Bedding was transferred daily for two weeks to ensure engraftment and after two additional weeks, behavioral analysis was performed on the ex-germ-free recipient mice. (**G**) The ex-germ-free mice who received the microbiome from stressed animals had increased time inactive by the tail suspension test, (**H**) no difference in the nestlet shred, and (**I**) decreased time in the open arms of the elevated plus maze in comparison with mice who received the microbiome from the naïve donors (Feale mice; n=6-7 mice/group; N= 1 experimental replicate; Student’s T tests).

### Microbiota from stressed animals can suppress systemic IFNγ production

To understand how microbiota from stressed mice may initiate anxiety- and depressive-like behaviors in germ-free mice, we first performed untargeted metabolomics on the serum from recipients. This revealed higher levels of stress hormones (**Figure 2A**) than the naïve microbiome recipients as expected. Numerous changes in other metabolic pathways were also evident, however, no trends detected. We next aimed to ensure that the different microbial communities didn’t cause changes in intestinal permeability. We first analyzed the gut physiology and function. The expression of several ion channels were analyzed as markers for proper digestion and function and no differences were found between the two groups (**Figure 2B**)[35,36]. Since germ-free mice have been reported to present with disturbed gut barrier integrity, the next step was examining markers of barrier junctions by qPCR and again, no differences were found (**Figure 2C**)[37]. Together, we conclude that there are no gross disruptions to gut permeability and stability.

**Figure 2:**
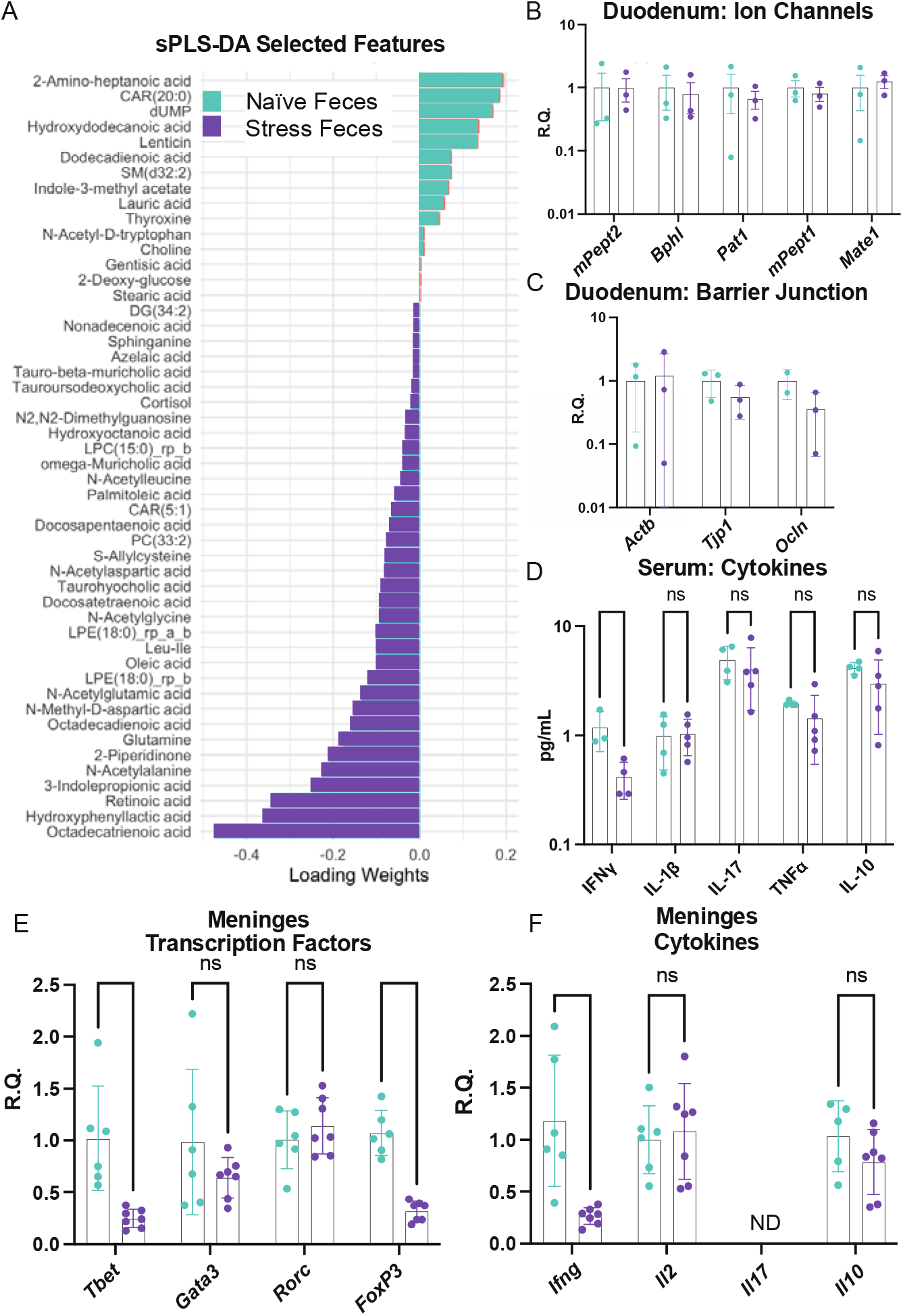
Ex-germ-free mice who received microbiome from stressed mice have lower Th1-associated signature. (**A**) Untargeted metabolomics performed on the serum of ex-germ-free mice who were colonized with the gut microbiota from either naïve or stressed donors. Data presented represent the significantly different annotated molecules by concentration as established by partial least squares analysis (Female mice; n=4-5 mice/group; N= 1 experimental replicate). qPCR on the duodenum of ex-germ-free mice targeting (**B**) ion channels and (**C**) epithelial tight junction associated transcripts (Female mice; n=2-3 mice/group; N= 1 experimental replicate; unpaired T tests corrected for multiple comparisons using the Benjamini, Krieger, and Yekutieli two-stage linear step up procedure). (**D**) Luminex analysis was performed on the serum from ex-germ-free mice. (Female mice; n=3-5 mice/group; N= 1 experimental replicate; unpaired T tests corrected for multiple comparisons using the Benjamini, Krieger, and Yekutieli two-stage linear step up procedure). Examination of the dural meninges from ex-germ-free mice by qPCR on (**E**) transcription factors associated with four types of CD4+ T helper cells and (**F**) their associated cytokines (Female mice; n=5-7 mice/group; N= 1 experimental replicate; unpaired T tests corrected for multiple comparisons using the Benjamini, Krieger, and Yekutieli two-stage linear step up procedure).

In recent years, the adaptive immune system and in particular, tissue resident CD4+ T cells, have emerged as a key player responsible for communication between the gut and brain in stress[38–41]. We performed Luminex analysis to quantify the concentrations of cytokines in the serum. The only cytokine that changed was IFNγ which was reduced in recipients of the stressed microbiome (**Figure 2D**). Next, select immune markers in the meninges, a site recently shown to host numerous immune cells and influence behaviors, were probed [42]. A decrease in the expression of *Tbet* in dural meninges, the transcriptional factor most associated with Type 1 anti-viral adaptive immunity and IFNγ expression, was noted. There was also a reduction *FoxP3*, the T regulatory transcription factor, by qPCR (**Figure 2E**). Expression of transcription factors linked to Type 2 (*Gata3*) or Type 17 immunity (*Rorc*) were not altered (**Figure 2E**). By analyzing expression of the signature cytokines of each of the T cell subsets, we found that only *Ifng* was decreased, paralleling our results by serum multiplex analysis (**Figure 2F**). Taken together, mice receiving the microbiome of stressed animals presented with a reduction in markers associated with Type 1 immunity, especially IFNγ which is a cytokine that has recently been linked to sociability by acting directly on neurons [42].

### Mice without *Lactobacillus* have fewer Th1 cells and are more susceptible to stress

So far, these data suggest that stress leads to a decrease in *Lactobacillus*. We have also shown that the transfer of microbiome from stressed animals was sufficient to drive anxiety and depressive-like behaviors and a decrease in IFNγ. To explore whether *Lactobacillus* was responsible for the changed IFNγ production, ASF animals were used. ASF mice have a defined gut composition consisting of eight commensal bacteria including two strains of *Lactobacillus* [18]. These gnotobiotic mice allow for the examination of a more complex gut microbiome than germ-free mice. These mice have a digestive and immune system that are closer to an SPF mouse than their germ-free counterparts [18]. To probe our hypothesis that *Lactobacillus* is responsible for IFNγ changes, gnotobiotic mice that have the complete ASF or ASF lacking the two species of *Lactobacillus*-*L. intestinalis* and *L. murinus* were generated. From here, they will be referred to as ASF (+L) and ASF (-L), respectively. These mice were reared in colonies that perpetuated these two gut microbial compositions. By flow cytometry, ASF(+L) and ASF(-L) mice have no differences in the number or percent of CD8+ T cells or CD4+ T cells (**Figure 3A, B, Supplement 1**). They also have no differences in the number of CD11b+ or B220+ cells indicating that there are no gross differences in monocyte or B cell populations (**Supplement 1**). However, of the CD4+ T cells there is a lower proportion of cells that are Tbet+ in the small intestine (**Figure 3C**). These data support the hypothesis that *Lactobacillus* are a novel regulator of Tbet expression and perhaps Type 1 immune responses. To further characterize these cells, we found that a lower proportion of the CD4+Tbet+ cells in ASF(-L) express CD69, a marker for activation and memory (**Figure 3D**).

**Figure 3:**
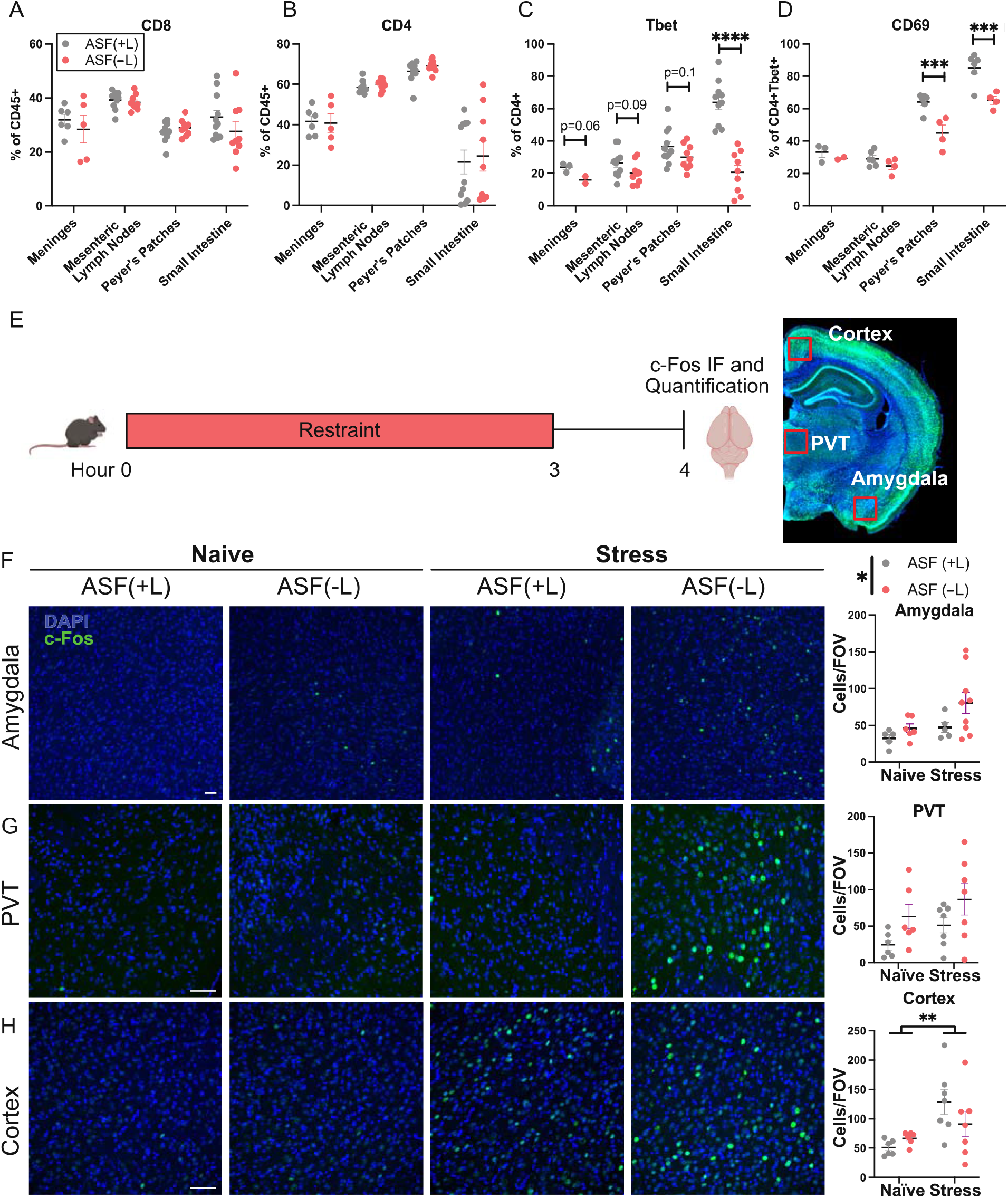
Mice without *Lactobacillus* from birth mirror immune and behavioral phenotypes of stressed mice. Flow cytometry analysis of dural meninges, mesenteric lymph nodes, Peyer’s patches, and lamina propria of the small intestine in either ASF (+L) or ASF(-L) mice. Cells gated on singlets, live cells, CD45+ and (**A**) CD8 or (**B**) CD4. CD4+ cells were further gated on (**C**) Tbet followed by (**D**) CD69. (Male and female mice; dural meninges from 2-3 mice were pooled leaving n=3-5 biological replicates. CD4, CD8, and Tbet had n=9-10 mice/group and N=2 experimental replicates; CD69 had n=4-5mice/group and N=1 experimental replicate; Two-way ANOVA followed by Sidak’s multiple comparison test with a single pooled variance). (**E**) Schematic of experimental design for F-H. After three hours of restraint c-Fos expressing cells were counted in the (**F**) Amygdala, (**G**) periventricular nucleus of the thalamus, and (**H**) Cortex. (Male and female mice; n=6-7 mice/group; N= 2 experimental replicates; Two-way ANOVA followed by Sidak’s multiple comparison test with a single pooled variance). Scale bars represent 30*μ*m.

To see if the presence or absence of *Lactobacillus* would alter the stress response, ASF(+L) and ASF(-L) mice were exposed to a mild acute stressor-restraint. We then performed c-Fos staining and quantification in areas of the brain involved in the stress response (**Figure 3D**). After a single restraint, ASF(-L) mice had higher activation in the stress response regions of the amygdala and the periventricular nucleus of the thalamus (PVT) than the ASF(+L) mice (**Figure 3F, G**). Notably, these trends were maintained in both the naïve and stressed mice. In the cortex, increased c-Fos expression was observed after stress in both groups, but no difference between groups (**Figure 3H**). These data show that *Lactobacillus* can influence Th1 cells and neuronal activation after acute stress.

### Mice lacking *Lactobacillus* have amplified behavioral responses to subclinical stress

While these data suggest that mice lacking *Lactobacillus* present with higher stress-induced neuronal activity as determined by c-Fos expression, it may not translate into long term behavioral changes. To test if absence of *Lactobacillus* leads to a change in depression- and anxiety-like behaviors, we developed a stress paradigm adapted from the unpredictable chronic mild stress paradigm previously used. This subclinical chronic stressor only uses restraint based on space constraints in the gnotobiotic isolators. For this stress paradigm, mice were restrained for 2 hours daily for 7 days (**Figure 4A**). Behavioral changes using this paradigm in SPF only appear after three weeks. There were no differences in anxiety or depression measures at baseline, however, after the subclinical stressor, only mice without *Lactobacillus* exhibited anxiety-like behavior as measured by the nestlet shred (**Figure 4B, Supplement 2**). The tail suspension test, a measure of depression-like behavior, also showed that the mice without *Lactobacillus* responded to the mild stressor (**Figure 4C**). A third measure of anxiety-like behavior, the elevated plus maze, revealed that both groups spent less time in the open arms after stress, though this may be due to habituation following multiple testing as other anxiety-like tests did not corroborate these results (**Figure 4D**). Together, these data indicate that mice lacking *Lactobacillus* in a simplified gnotobiotic microbiome are more susceptible to stress when compared to similar microbiomes that contain *L. intestinalis* and *L. murinus*.

**Figure 4:**
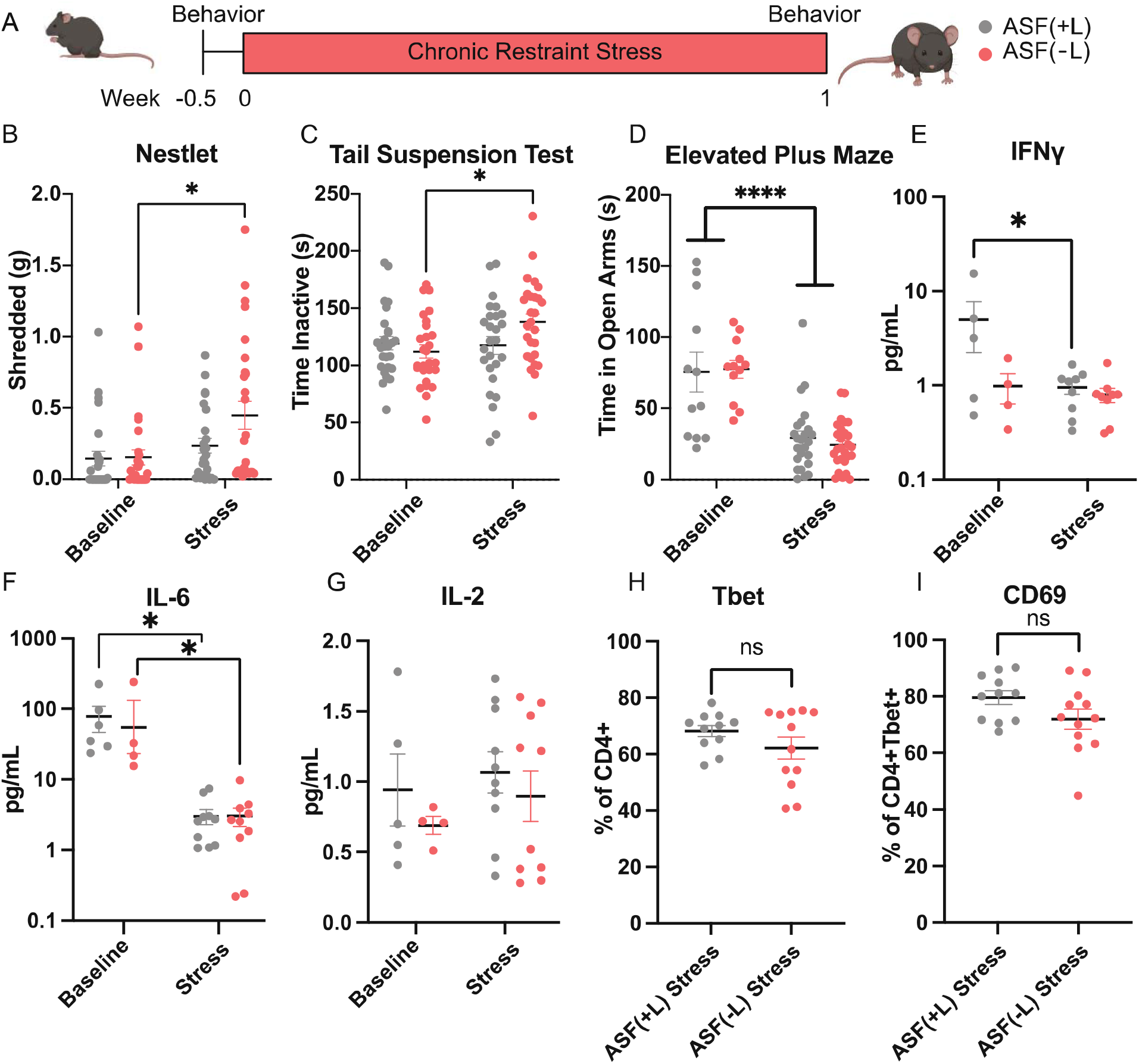
Mice without *Lactobacillus* are more behaviorally susceptible to subclinical stressors than mice with *Lactobacillus*. (**A**) Schematic representing the modified subclinical stressor aimed at preventing bacterial contamination. ASF(+L) and ASF(-L) were restrained for two hours daily for one week in sterile tubes flanked by behavior testing. Behavioral testing at baseline and after stress included (**B**) the nestlet shred, (**C**) tail suspension test, and (**D**) the elevated plus maze (Male and female mice; n=24-26 mice/group; N=3 experimental replicates; One-way ANOVA followed by Dunnett’s multiple comparison test with a single pooled variance). (**E**) Luminex analysis on the serum of ASF(+L) and ASF(-L) mice before and after stress (Male mice; n=4-10 mice/group; N=2 experimental replicates; One-way ANOVA followed by Dunnett’s multiple comparison test with a single pooled variance). IFNγ, (**F**) IL-6, and (**G**) IL-2 are shown. (**H**) Flow cytometric analysis of the lamina propria of the small intestine in stressed ASF(+L) and ASF(-L) mice gated on singlets, live cells, CD45+, CD4+, and Tbet+ followed by (**I**) CD69 (Male and female mice; n=11-12 mice/group; N=2 experimental replicates; Student’s T tests).

To determine whether the serum concentrations of IFNγ followed what was observed in mice colonized with the gut microbiota from stressed mice (**Figure 2E**), serum levels of cytokine were tested before and after stress. As hypothesized, ASF(-L) mice have lower concentration of serum IFNγ than ASF(+L) mice (**Figure 4E**). After stress, serum IFNγ decreases only in the ASF(+L) mice. These data indicate that a reduction in IFNγ (and thus a higher homeostatic concentration of IFNγ) may be necessary for psychological resilience to environmental stressors. To confirm that the concentrations of IFNγ were not simply a result of overall changes in the pro-inflammatory environment, we also measured other cytokines associated with environmental stressors-IL-6 and IL-2 [43,44]. Neither are reliant upon the presence of *Lactobacillus* indicating that this is likely an IFNγ specific effect (**Figure 4F, G**). In further support of the hypothesis that Type 1 immunity is modulated during stress, we also observe that the intestinal differences in the proportion of Tbet+ and Tbet+CD69+ cells between ASF(+L) and ASF(-L) mice at baseline (**Figure 3C, D**) equilibrate after stress (**Figure 4H, I**). Overall, by utilizing ASF mice, we have shown that systemic IFNγ is dependent on the presence of *Lactobacillus* in the gut microbiota and that in the absence of *Lactobacillus* and IFNγ mice are more susceptible to environmental stress.

### Circulating IFNγ may reduce stress susceptibility

Other groups have previously shown that genetic knockout of IFNγ increases anxiety-like behavior, but little work has focused on more physiological systems [45]. In order to determine a causative relationship between circulating IFNγ and stress responses, either IFNγ or anti-IFNγ neutralizing antibody were administered to animals exposed to the acute stress paradigm (**Figure 5A, Supplement 3**). In the amygdala, there was a significant increase in c-Fos expression between stress animals compared to naïve animals (**Figure 5B**). Animals given IFNγ i.p. before stress appeared to have lower c-Fos expression after the acute stressor than controls, while those given the neutralizing antibody seemed to have heightened responses to the acute stressor both in the amygdala and the periventricular nucleus of the thalamus (**Figure 5B-C**). The cortex showed similar patterns to the manipulation of *Lactobacillus* (**Figure 5H**)-there was an increase in c-Fos expression in response to stress, but no difference based on manipulation of IFNγ (**Figure 5D**). Overall, these data support previous work that circulating IFNγ can regulate behavioral responses and may be produced in response to the gut microbiota[45].

**Figure 5:**
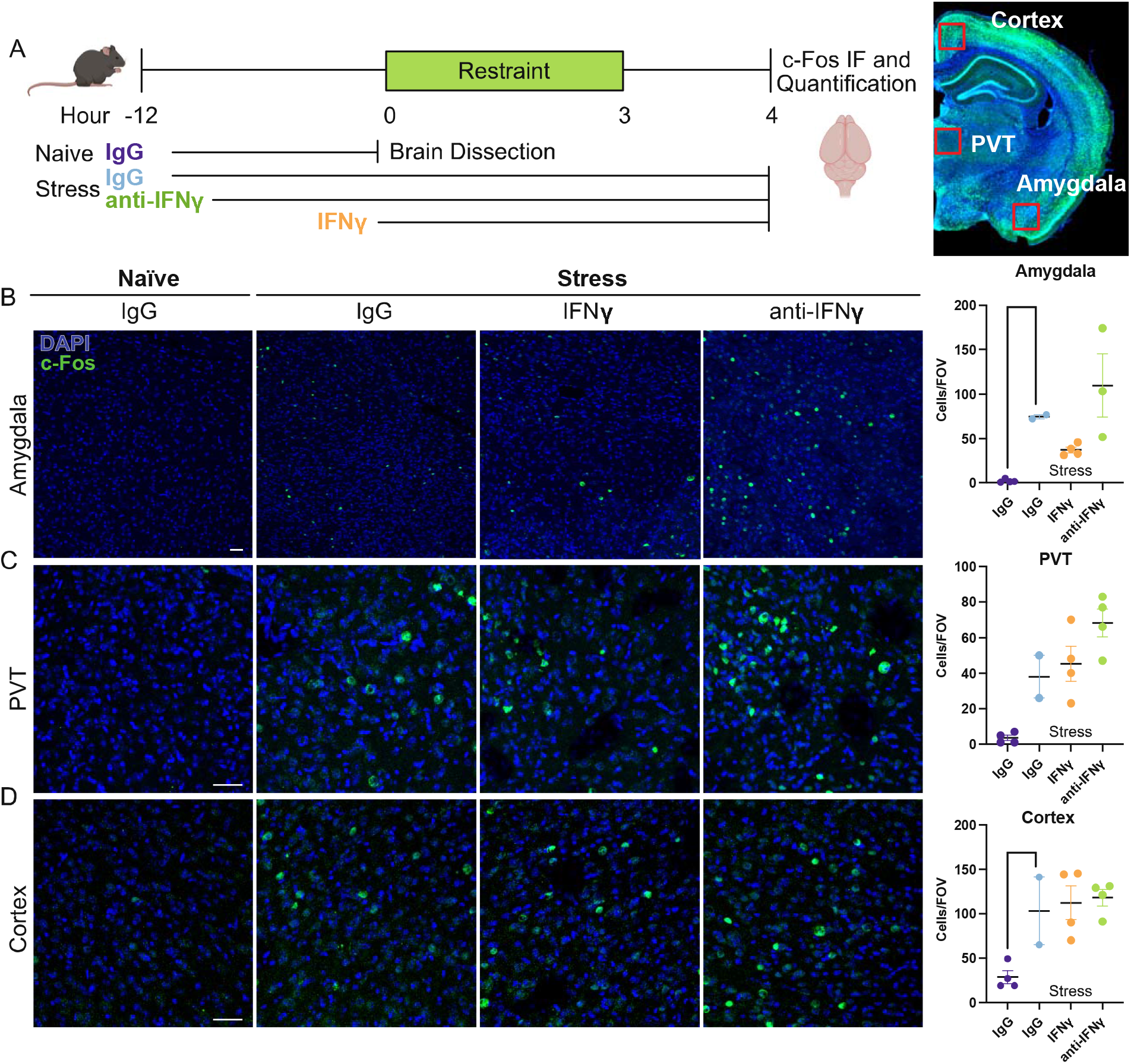
Circulating IFNγ may drive susceptibility to stress. (**A**) Schematic of experimental design. Mice were treated i.p. with anti-INFγ neutralizing antibody or IgG 12 hours before a three-hour acute restraint stress. One group was given recombinant IFNγ immediately before acute restraint stress. Brains were dissected for c-Fos quantification before or after stress. (**B**) c-Fos quantification in the amygdala, (**C**) periventricular nucleus of the thalamus, and (**D**) cortex (Male mice; n=2-4/group; N= 1 experimental replicate; One-way ANOVA followed by Dunnett’s multiple comparison test with a single pooled variance). Scale bars represent 30*μ*m.

## 3.2 Discussion

Historically, researchers have used blunt tools in order to study the gut microbiota. Germ-free rodents were developed in the early 1940s and can be reconstituted with the microorganisms of interest; however, there are significant challenges in interpreting and extrapolating results from these studies [46]. They are immunocompromised, have high levels of anxiety-like behavior, and do not respond to novel microorganisms in the same way a specific pathogen free (SPF) mouse would [37,47]. Another common method to study commensal bacteria is to disrupt the microbiome of SPF mice using antibiotics and colonizing with the microorganism of interest. However this comes with other issues: antibiotic use can cause off target effects and the engraftment of the target microbe is not stable [48,49]. Due to these challenges, it has been difficult to understand the mechanistic relationship between *Lactobacillus* residing in the gut microenvironment and mood thus far.

Here, we have shown that the gut microbiome is changed in mice subjected to environmental stress and that this new population is sufficient to drive anxiety- and depressive-like behavior when transferred to germ-free mice. Mice raised with a known consortium of gut bacteria that contains *Lactobacillus* are more resistant to both acute stressor and more chronic mild stressors than mice without *Lactobacillus*. This difference seems to be IFNγ-dependent and likely results from an indirect exposure of Th1 cells to bacterial components.

While the exact mechanism of Th1 sensing of *Lactobacillus* is unclear, we hypothesize that the communication is occurring through bacterial extracellular vesicles (EVs) [50]. There is evidence that EVs from *Lactobacillus* administered in vivo can reduce environmental stress responses [51]. Beyond EVs, there is likely increased Th1 activation and IFNγ production occurring through canonical intestinal immune development through sampling by M cells along with antigen presentation by dendritic cells, or interaction with small molecules excreted by the bacteria [52,53].

Our work supports previous work implicating IFNγ in the stress response and behavior [45]. Correlations between stress, depression, and IFNγ have previously been shown in the clinic, but the causative relationship and mechanism have not been established [54,55]. Recent studies have also shown that meningeal sources of IFNγ can act directly on neuronal IFNγ receptors to promote the normal development of social behavior[42]. In addition to neurons, astrocytes and premyelinating oligodendrocytes also have IFNγ receptors, indicating that there may be several responding cell types in the brain [56,57]. Overall, there are likely both direct and indirect mechanisms of IFNγ influence on neuronal circuits and behavior.

These results provide a novel framework to understand the roles of the bacterial gut microbiome and the immune system in mood disorders. Innovative methods were used to show that *Lactobacillus* are indeed driving IFNγ production - likely via antigen presentation and not due to direct pattern recognition receptor activation. We have also shown that the IFNγ produced in response to *Lactobacillus* is sufficient to provide resilience in response to environmental stressors, as had been previously observed in clinical settings. This work is a key step for understanding the role of homeostatic physiology in developing psychological resilience and may help to direct the search for probiotic supplements that may operate as supplemental therapies for mood disorders.

## Supporting information

statistics

