## Supplementary material for "*Lactobacillus* maintains IFNγ homeostasis to promote behavioral stress resilience": Suplemental figures

**Supplement 1: Phenotyping of ASF(+L) and ASF(-L) by flow cytometry.** (**A**) Gating strategy for flow cytometric analysis of ASF(+L) and ASF(-L) mice. Representative plot of Peyer’s patch cells**.** (**B**) Number of B220+ B cells (**C**) CD11b+ monocytes and (**D**) TCRb+ T cells. (**E**) Percent of CD4+ T cells that are RORgt+. (Male and female mice; dural meninges from 2-3 mice were pooled leaving n=3-5 biological replicates. Mesenteric lymph nodes, Peyer’s patches, and lamina propria of the small intestine had n=9-10 mice/group and N=2 experimental replicates; CD11b had n=3-5/group and N=1 experimental replicate. Two-way ANOVA followed by Sidak’s multiple comparison test with a single pooled variance).

**Supplement 2: ASF(+L) and ASF(-L) do not show behavioral differences in the open field test or marble burying test** (**A**) 16S sequencing of fecal pellets from ASF(+L) and ASF(-L) mice. (**B**) Open field test time in center and (**C**) total distance traveled before and after stress. (**D**) Marble burying test after stress. (Male and female mice; n=24-26 mice/group; N=3 experimental replicates; One-way ANOVA followed by Dunnett’s multiple comparison test with a single pooled variance).

**Supplement 3: c-Fos expression in CA1 region of the hippocampus in ASF(+L) and ASF(-L) mice and mice treated with IFNγ.** (**A**) After three hours of restraint c-Fos expressing cells were counted in the CA1 region of the hippocampus (Male and female mice; n=6-7 mice/group; N= 2 experimental replicates; Two-way ANOVA followed by Sidak’s multiple comparison test with a single pooled variance). (**B**) Mice were treated i.p. with anti-INFγ neutralizing antibody or IgG 12 hours before a three-hour acute restraint stress. One group was given recombinant IFNγ immediately before acute restraint stress. Brains were dissected for c-Fos quantification before or after stress. c-Fos quantification in the CA1 region of the hippocampus (Male mice; n=2-4/group; N= 1 experimental replicate; One-way ANOVA followed by Dunnett’s multiple comparison test with a single pooled variance).
